# Characterizing the SASP-Dependent Paracrine Spreading of Senescence Between Human Brain Cell Types

**DOI:** 10.64898/2026.02.10.705129

**Authors:** Taylor Russo, Markus Riessland

**Author notes:** **Correspondence:** Markus Riessland.

## Abstract

One of the defining phenotypes of a senescent cell is the senescence-associated secretory phenotype (SASP), which can propagate senescence in neighboring cells both *in vitro* and *in vivo*. Importantly, this paracrine spreading of senescence can act in a cell non-autonomous manner, influencing neighboring cell populations and contributing to immune cell recruitment. As cellular senescence has recently been linked to both age-related neurodegenerative phenotypes and local inflammation and is more clearly defined across brain cell types in a cell-type-dependent manner, an urgent question remains regarding how a cell-type-specific paracrine spreading of senescence occurs in the brain. Here, we set out to profile the cell-type-specific features of the SASP and characterize the directionality of paracrine senescence-spreading between major brain cell types. Through this analysis, we identified key SASP ligand-receptor pairs involved in this paracrine dissemination. Targeting these factors with specific inhibitors, we prevented the paracrine spreading of senescence in a brain cell-type-dependent manner. Taken together, we identified specific SASP targets for therapeutic intervention in the context of human brain cells and thereby informed the SASP-dependent reaction of immune cells and age-related tissue dysfunction across both normal aging and models of neurodegenerative disease.

## Introduction

Cellular senescence is a complex stress response induced by a variety of triggers and is characterized by key hallmarks, including the senescence-associated secretory phenotype (SASP) (*1–3*). Senescent cells accumulate across multiple tissues with age and have been shown to contribute to various neurodegenerative phenotypes (*4–6*). Senescent cells have been reported to display both beneficial, such as promoting tissue repair, and harmful effects, including contributing to chronic inflammation and tissue dysfunction (*7*). Despite these dual roles, recent studies have highlighted their contribution to functional decline as well as inflammatory phenotypes in neurodegeneration and brain aging points towards a need for a better understanding of their biology to potentially mitigate associated damage (*8, 9*). Specifically, the SASP has been shown to induce a paracrine spreading of senescence to neighboring non-senescent cells (*10*). The initiating senescent cells, which first undergo the senescence program in response to stress, are commonly referred to as having a ‘primary senescence’ phenotype, whereas neighboring cells that subsequently acquire a senescence phenotype through exposure to SASP factors develop a distinct molecular signature which has been termed ‘secondary senescence’ (*11*). Importantly, there have been recent reports of transcriptional heterogeneity across senescent cells (*12*), as well as our recent profiling and characterization of cell-type-specific profiles of senescence across various human brain cell types demonstrating their molecular diversity (*13*). In addition to cell type, the inducing stressor as well as the tissue in which the senescent cells reside are factors contributing to heterogeneity of senescence hallmarks (*14*). Although no single feature ultimately defines a senescent cell, the SASP is a well-characterized phenotype that promotes immune cell recruitment and drives a paracrine spreading of senescence, contributing to age-related decline across multiple tissues (*10*).

Recent advancements aimed to delineate the mechanisms of the paracrine spreading of senescence both *in vitro* and *in vivo* have shown that the accumulation of reactive oxygen species (ROS) and the NFκ-B pathway are key mediators (*15*). Specifically, senescence-associated mitochondrial dysfunction (SAMD) has been linked to ROS accumulation and subsequent NFκ-B pathway modulation of the SASP leading to DNA damage in bystander cells (*16*). Additional work *in vivo* has highlighted IL1-β as a key mediator of primary versus secondary senescence, which is also both tissue- and inducer-dependent (*14*). Interestingly, *in vivo* models utilizing the transplantation of senescent cells into young mice have demonstrated a spreading of senescence to both local and distant host tissue via the SASP (*17*). Other key SASP factors include TGFβ, IGF1, IL-6, IL-8, and CCL2, where conditioned media (CM) experiments were used to delineate which SASP factors are critical in mediating the paracrine transmission of senescence between neighboring cells (*18*). The use of a TGFβR1 antagonist was shown to specifically reduce paracrine spreading of senescence, as opposed to an autocrine spread (*19*). Importantly, it has been shown that paracrine spreading of senescence can occur between different cell types and is capable of producing a stable secondary senescence phenotype in the receiving cells (*19*). Additionally, it is well established that different cell types express different SASP profiles (*3, 20*). Overall, the spread of senescence clearly contributes to age-related tissue dysfunction (*6*). However, in the context of brain cell senescence, it remains unclear which cell types are the main mediators, which are the most susceptible to entering secondary senescence, and which key SASP factors (e.g. ligand-receptor pairs) may mediate this potentially cell-type-specific paracrine spread of senescence. A better understanding of how spreading of senescence between neurons, astrocytes, microglia, oligodendrocytes and endothelial cells may contribute to neurodegenerative phenotypes, immune cell activation, and ultimately a loss of blood-brain-barrier integrity (*2*) would elucidate a targetable mechanism of age-related disease.

Here, we sought to unravel the relationship between key brain cell types (astrocytes, endothelial cells, microglia, oligodendrocytes, and neurons) in the context of a paracrine spreading of senescence via the SASP. We utilized our previously established *in vitro* DNA damage-induced human brain cell line senescence model (*13*) and CM experiments to profile the cell-type-dependent SASP, characterize the directionality of a paracrine spreading of senescence between the relevant cell types, identify key SASP ligands and receptors that mediate the cell-type-specific spread, and target these factors using various inhibitors in an attempt to prevent the paracrine spreading of senescence. We demonstrate that a cell-type-specific SASP profile of each brain cell type drives differential induction of secondary senescence, where some cell types can induce senescence in themselves as well as in other cell types, while other cell types are only capable of receiving secondary senescence induction, but cannot spread. Importantly, we identified both cell-type-specific and common SASP ligands and receptors, which we successfully targeted to prevent the induction of secondary senescence depending on the cell types communicating with one another. Taken together, this work gives key insights into the mechanisms of paracrine spreading of senescence between brain cell types *in vitro* and offers potential therapeutic targets to prevent this spreading, which may in turn help to alleviate age-related tissue decline and inflammaging.

## Results

### Spreading of senescence in human brain cell lines

In order to expand upon our previous characterization of 5-Bromodeoxyuridine (BrdU) induced senescent human brain cell types (*13*), we set out to elucidate SASP-specific alterations across the different human cell lines, which may drive differential responses in terms of a spreading phenotype. To do so, we utilized conditioned media (CM) from control (DMSO-treated) and senescent (100 µM BrdU-treated) human brain cell types including endothelial cells (pink), microglia (yellow), oligodendrocytes (green), astrocytes (purple), and neurons (blue) (Figures 1A,B). First, CM samples from each cell line were profiled by nucleobase Enabled Localized Immunoassay with Spectral Addressing (nELISA) technology using a Core Immune panel of 286 cytokine factors. Assessing alterations of these factors across the non-senescent (DMSO-treated) and senescent (BrdU-treated) groups, we observed cell-type-specific upregulation of distinct SASP components, revealing unique profiles in senescent endothelial cells, microglia, oligodendrocytes, astrocytes, and neurons (Figure 1C). These results revealed that while some cytokines were commonly upregulated across senescent brain cell types, others showed cell-type-specific increases, consistent with previous reports that SASP profiles vary depending on cellular context (*3, 20*).

**Figure 1:**
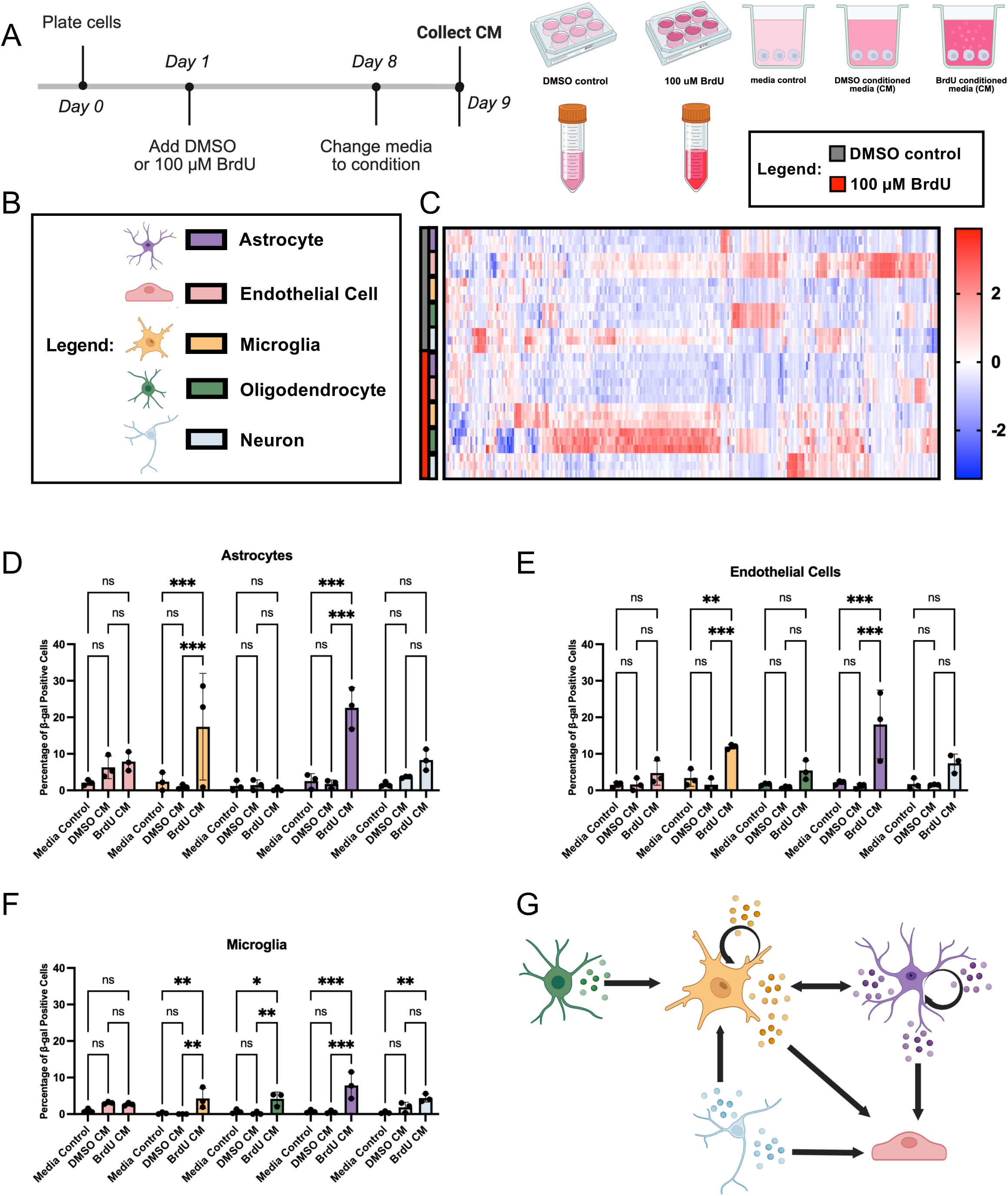
Spreading of senescence in human brain cell lines. A) 9-day timeline of conditioned media (CM) collection from human cell lines following treatment with DMSO control or 100 µM BrdU (created with BioRender). Schematics of DMSO and BrdU treatment to collect CM followed by CM treatment (including media control) on non-senescent cells (created with BioRender). B) Legend for cell type images and colors for data shown in (C-G) where endothelial cells are pink, microglia are yellow, oligodendrocytes are green, astrocytes are purple, and neurons are blue. C) Z-score heatmap showing Core Immune panel (Nomic Bio) cytokine expression (including SASP factors) in all cell types based on nELISA analysis of CM from DMSO (grey) and BrdU (red) treated cell lines (n=3 replicates). D) Quantification of percentage of SA β-gal positive senescent endothelial cells (pink), microglia (yellow), oligodendrocytes (green), astrocytes (purple), and neurons (blue) following 7-day treatment with astrocyte DMSO and BrdU CM as well as a cell-type-dependent media control (n=3 replicates). E) Quantification of percentage of SA β-gal positive senescent endothelial cells (pink), microglia (yellow), oligodendrocytes (green), astrocytes (purple), and neurons (blue) following 7-day treatment with endothelial cell DMSO and BrdU CM as well as a cell-type-dependent media control (n=3 replicates). F) Quantification of percentage of SA β-gal positive senescent endothelial cells (pink), microglia (yellow), oligodendrocytes (green), astrocytes (purple), and neurons (blue) following 7-day treatment with microglia DMSO and BrdU CM as well as a cell-type-dependent media control (n=3 replicates). G) Schematic depicting results from (D-G) showing which cell types were able to induce senescence (significantly increased percentage of SA β-gal positive cells) in which other cell types. Data was analyzed by two-way ANOVA with Tukey’s multiple comparisons test (D-F). All graphs show mean with error bars depicting standard deviation (ns, p>0.05, * p<0.05, ** p<0.01, *** p<0.001).

We next sought to determine which of these uniquely upregulated SASP factors drive the spread of senescence between different brain cell types. Hence, we treated each of the five human cell lines with non-senescent (DMSO, grey) CM and senescent (BrdU, red) CM from every other cell type (i.e. astrocytes treated with DMSO CM and BrdU CM from microglia). We also included an unconditioned media control group to ensure that culturing one cell line in another cell’s media had no effect on senescence hallmark expression (Figure 1A). To analyze the induction of senescence, we quantified the percentage of senescence-associated β-gal (SA β-gal) positive cells in each treatment group, building on our prior characterization showing that BrdU treatment significantly increases SA β-gal staining across all five human brain cell types (*13*). We observed a significant increase in SA β-gal positive astrocytes and endothelial cells when they were treated with BrdU CM from microglia or from other astrocytes (Figure 1D, E). Concurrently, microglia demonstrated a significant increase in SA β-gal following treatment with BrdU CM from other microglia, oligodendrocytes, astrocytes, or from neurons (Figure 1F). In contrast, both neurons and oligodendrocytes showed no significant change in the number of SA β-gal positive senescent cells when treated with BrdU CM from any cell type (data not shown). This suggests that these cell types are less susceptible to paracrine induction of secondary senescence.

Overall, we demonstrated that endothelial cells were susceptible to SASP-mediated secondary senescence but were unable to induce senescence in any other cell type (primary senescence) – even in other endothelial cells (Figure 1G). Most strikingly, we observed that astrocytes and microglia appeared to be the main drivers of senescence spread, where they expressed the strongest primary senescence phenotype and were both able to induce senescence in endothelial cells, onto themselves, and onto each other (Figure 1G). Taken together, these data demonstrate that each senescent human brain cell type displays a unique SASP profile that differentially drives paracrine spreading of senescence in a cell-type-specific manner. Furthermore, each cell type exhibits a distinct response to SASP-exposure. Importantly, astrocytes and microglia appear to be key primary senescent cell types capable of driving a secondary senescence phenotype in other relevant human brain cells.

### Astrocytes and microglia as the primary drivers of secondary senescence

Based on our SA β-gal results demonstrating that astrocytes and microglia appear to be the primary drivers capable of inducing senescence in many brain cell types (Figure 1G), we set out to further characterize the secondary senescence phenotype that their SASP induced in responding cells. To do so, we utilized astrocyte- and microglia-derived CM and tested combinations in which senescence was spread to the same cell type (astrocyte→astrocyte and microglia→microglia) as well as between these two cell types (astrocyte→microglia and microglia→astrocyte). Additionally, since we did not see a sufficient secondary senescence response from endothelial cell CM (Figure 1), we utilized this CM as a control group. We paneled the expression of senescence markers *CDKN1A* (p21), *CDKN2A* (p16), *CDKN2C* (p19), and *LMNB1*, guided by our previous characterization demonstrating modulation of these factors following BrdU treatment in the five human brain cell types (*13*). When astrocytes were treated with DMSO (grey) and BrdU (red) CM from the three brain cell types, we observed a significant upregulation of *CDKN1A* specifically following exposure to astrocyte-derived BrdU CM (Figure 2A). Astrocyte BrdU CM also induced significant increases in *CDKN2D* and *LMNB1* in astrocytes (Figure 2A), while *CDKN2A* levels remained unchanged. Additionally, microglia BrdU CM led to a significant upregulation of *CDKN2D* in astrocytes (Figure 2A). In contrast, treatment with BrdU CM from endothelial cells did not produce significant alterations in any senescence markers in astrocytes. Next, we treated microglia with DMSO and BrdU CM from the three brain cell types. Astrocyte-derived BrdU CM induced a significant downregulation of *CDKN1A* and *LMNB1* expression in microglia (Figure 2B), while microglia BrdU CM had no effect. Similarly to astrocytes, we observed no significant changes in senescence marker expression following treatment with endothelial cell BrdU CM (Figure 2B).

**Figure 2:**
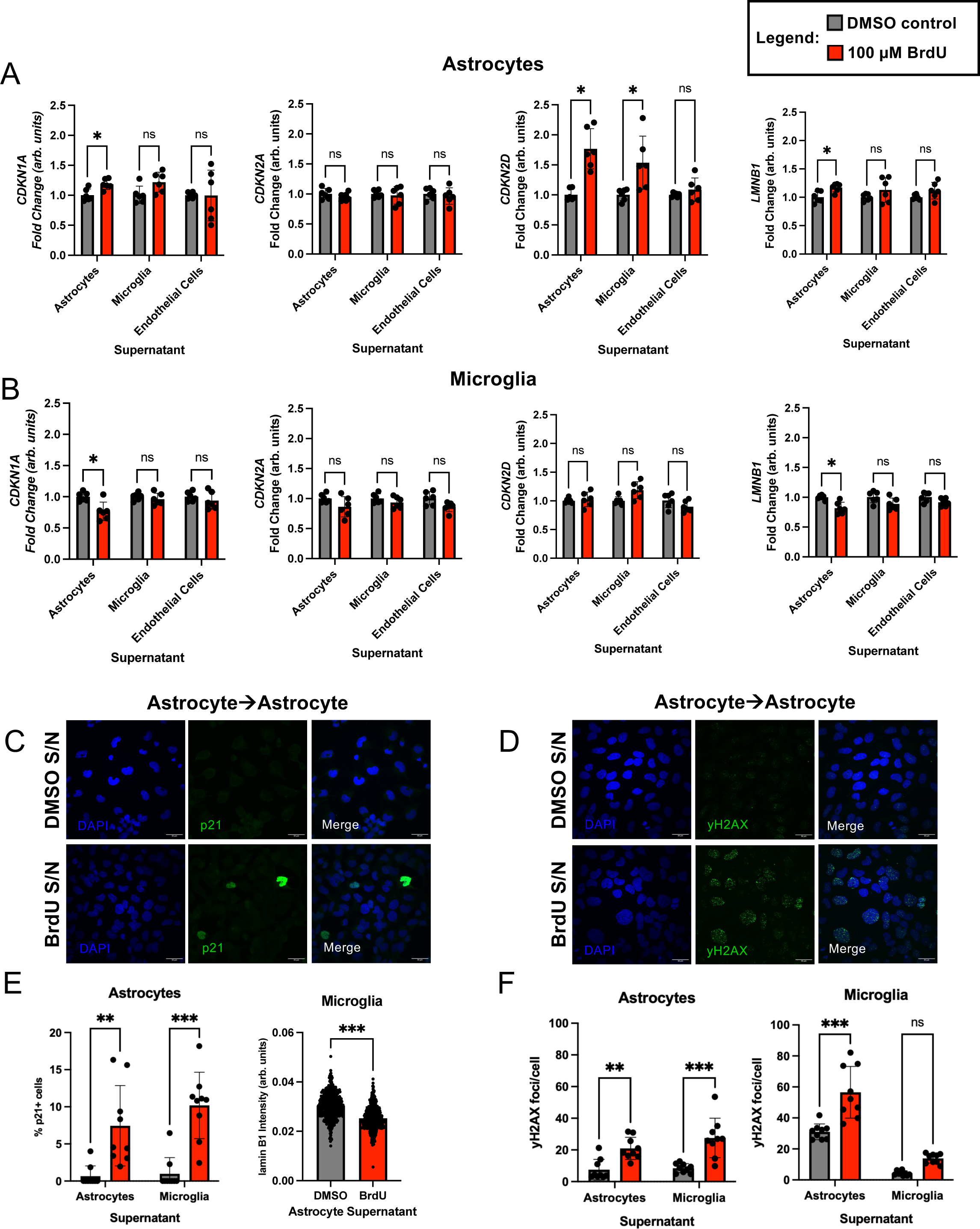
Astrocytes and microglia as the primary drivers of senescence phenotypes. A) Quantitative PCR of astrocyte, microglia, and endothelial cell DMSO CM (grey) and 100 µM BrdU CM (red) treated astrocytes for *CDKN1A*, *CDKN2A*, *CDKN2D*, and *LMNB1* (n=6 replicates). B) Quantitative PCR of astrocyte, microglia, and endothelial cell DMSO CM (grey) and 100 µM BrdU CM (red) treated microglia for *CDKN1A*, *CDKN2A*, *CDKN2D*, and *LMNB1* (n=6 replicates). C) Confocal images of astrocytes treated with DMSO CM and BrdU CM from astrocytes with staining for p21 (green) with DAPI (blue). D) Confocal images of astrocytes treated with DMSO CM and BrdU CM from astrocytes with staining for γH2AX (green) with DAPI (blue). E) Quantification of percentage (%) of p21 positive cells in astrocytes treated with DMSO CM and BrdU CM from astrocytes and microglia (n=9 replicates). Quantification of lamin B1 intensity in microglia treated with DMSO CM and BrdU CM from astrocytes (n=9 replicates). F) Quantification of the number of γH2AX foci per cell in astrocytes treated with DMSO CM and BrdU CM from astrocytes and microglia (n=9 replicates). Quantification of the number of γH2AX foci per cell in microglia treated with DMSO CM and BrdU CM from astrocytes and microglia (n=9 replicates). All graphs show mean with error bars depicting standard deviation (ns, p>0.05, * p<0.05, ** p<0.01, *** p<0.001). Data analyzed by unpaired two-tailed t-test (A-B), two-way ANOVA with Tukey’s multiple comparisons test (E, F), and unpaired Mann-Whitney t-test (E). All scale bars are 30 µm. *Note: a rendering error in the original preprint version caused a portion of the yH2AX DMSO S/N panel in D to display as partially blank black space, which has been corrected here to restore the complete original image data.*

To further panel senescence hallmarks following the various CM treatments, we utilized immunostaining (Figure S1). We observed that treatment with both astrocyte- and microglia-derived BrdU CM resulted in a significant increase in the percentage of p21-positive astrocytes (Figure 2C, E). In line with this, our quantification of p21 staining intensity showed a significant increase in astrocytes following treatment with BrdU CM from both astrocytes and microglia (Figure S2A). Subsequent staining in microglia with BrdU CM from astrocytes demonstrated a significant reduction in lamin B1 intensity, which is indicative of a senescence phenotype (Figure 2E). We also stained both astrocytes and microglia for γH2AX (pS139), a DNA damage marker associated with senescence-associated heterochromatin foci (SAHF) marker. In astrocytes, treatment with astrocyte-and microglia-derived BrdU CM led to a significant increase in the number of γH2AX foci per cell. In microglia, a similar increase was observed following astrocyte BrdU CM treatment (Figure 2D, F). Additional quantification of γH2AX intensity showed a significant increase in both astrocytes and microglia following treatment with both astrocyte and microglia BrdU CM (Figure S2B).

Lastly, given our previous work identifying mitochondrial and lysosomal dysfunction as hallmarks of BrdU-induced senescence in human cell lines (*13*), we investigated these phenotypes in CM-treated astrocytes and microglia. Using Mitotracker Red CMXRos staining, we observed a significant decrease in intensity (indicative of mitochondrial membrane potential) in both cell types following treatment with astrocyte-and microglia-derived BrdU CM (Figure S2C). Quantification of mitochondrial and lysosomal mass showed a significant decrease in the number of mitochondrial per cell in astrocytes following treatments with BrdU CM from both astrocytes and microglia (Figure S2D), with no significant changes in microglia with any CM treatment. Lysosomal mass remained unchanged in both cell types following treatment with astrocyte- or microglia-derived BrdU CM (Figure S2E). Taken together, this further characterization of senescence hallmarks following treatments with senescent CM from the primary senescent-driving cell types (astrocytes and microglia) indicates that they have the potential to induce different senescence hallmarks in the receiving secondary senescent cell types and that these changes are also dependent on the cell type that is being induced. Thus, the paracrine spreading of senescence is cell-type-specific both in the primary/sending cell and in the secondary/receiving cell.

### Multiplexed immunoassay analysis identifies distinct SASP signatures of astrocytes and microglia

After establishing the directionality of senescence spread among astrocytes, endothelial cells, microglia, oligodendrocytes, and neurons, and identifying the resulting cell-type-specific paracrine induction of senescence hallmarks in receiving cells, we revisited our nELISA data (Figure 1C) to further characterize the SASP profiles of the primary senescence drivers. To elucidate the specific factors upregulated in BrdU CM from each human cell line, we generated volcano plots to visualize significantly (p<0.05) upregulated (red, log2fc > 1) and downregulated (blue, log2fc < -1) cytokines. Astrocyte BrdU CM showed an upregulation of matrix metalloproteinases MMP-1 and MMP-3 along with a significant increase in GDF-15 (MIC-1) levels and CCL2 signal (Figure 3A). GDF-15 has been established as a stress response factor, but more recently has been discussed as a novel biomarker in human systemic senescence-related aging (*21*). CCL2 is a chemokine secreted by senescent cells which has been targeted therapeutically in the context of colorectal cancer progression (*22*). Downregulated factors in astrocyte BrdU CM included CXCL12 alpha and beta, which have been shown to be cleaved by DPP4 generating metabolites with altered bioactivity (*23*). Although CXCL12 is an established SASP factor, DPP4 has been shown to be elevated in senescent fibroblasts, potentially contributing to its processing (*24*). Endothelial cell BrdU CM revealed an upregulation of four specific factors including matrix metalloproteinases MMP-12 and MMP-1, along with uPA and EGF, which have both been shown to be involved in mediating a senescence phenotype (*25, 26*) (Figure 3B). Most interestingly, BrdU CM from endothelial cells displayed an overall widespread downregulation of immune factors including CCL2, CXCL12 alpha and beta, and MIF, which has been shown to protect cells from entering a state of senescence (*27*). BrdU microglia CM showed an upregulation of many factors, including CCL2, MIF (which has also been shown to regulate the expression of other cytokines (*28*)), CXCL12 beta, and MMP-1,3,12,13 and a downregulation of CXCL12 alpha (Figure 3C). Oligodendrocyte BrdU CM showed an upregulation of MMP-1, CCL2, CXCL12 alpha and beta, and IL-2 RA (Figure 3D). Lastly, neuronal BrdU CM displayed a downregulation of MIF and an upregulation of MMP-3, CCL2, CCL5, and CCL20 among many other cytokines (Figure 3E). Taken together, BrdU CM (red) from each human brain cell type displays a unique alteration in the 286 immune factors (Figure 3G), which may explain the directionality of the paracrine spreading of senescence we observed using SA β-gal analysis (Figure 1G).

**Figure 3:**
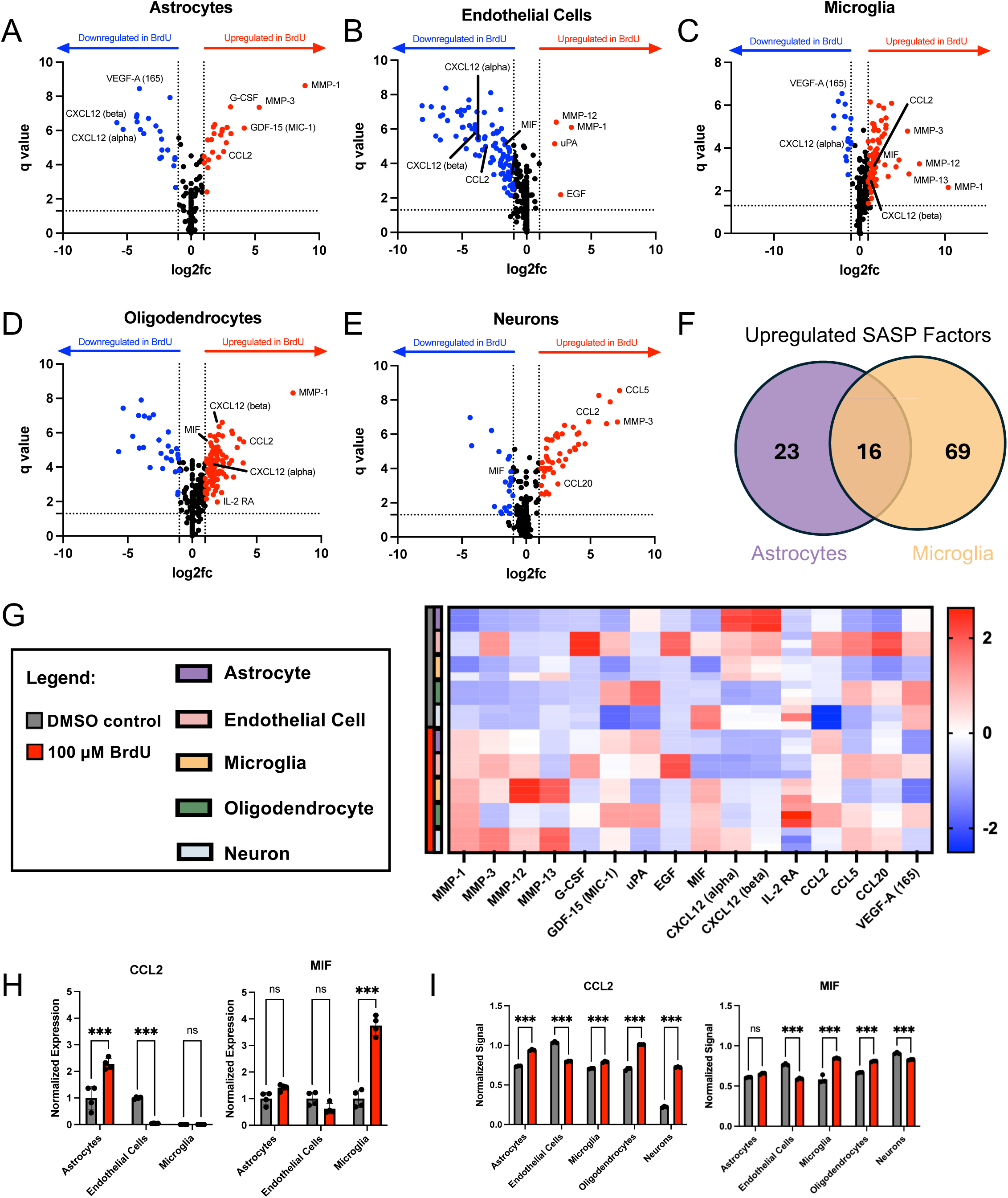
Multiplexed immunoassay analysis identifies distinct SASP signatures of astrocytes and microglia. A) Volcano plot of Nomic Bio Core Immune panel (268 cytokines) in DMSO and BrdU CM from astrocytes showing factors that are upregulated in BrdU CM treated (red) (log2fc > 1, p-value < 0.05) and downregulated in BrdU CM treated (blue) (log2fc < -1, p-value < 0.05). Black data points indicate factors that do not reach the threshold (−1 < log2fc < 1 | p-value > 0.05). B) Volcano plot of Nomic Bio Core Immune panel cytokines in DMSO and BrdU CM from endothelial cells showing factors that are upregulated in BrdU CM treated (red) (log2fc > 1, p-value < 0.05) and downregulated in BrdU CM treated (blue) (log2fc < -1, p-value < 0.05). Black data points indicate factors that do not reach the threshold (−1 < log2fc < 1 | p-value > 0.05). C) Volcano plot of Nomic Bio Core Immune panel cytokines in DMSO and BrdU CM from microglia showing factors that are upregulated in BrdU CM treated (red) (log2fc > 1, p-value < 0.05) and downregulated in BrdU CM treated (blue) (log2fc < -1, p-value < 0.05). Black data points indicate factors that do not reach the threshold (−1 < log2fc < 1 | p-value > 0.05). D) Volcano plot of Nomic Bio Core Immune panel cytokines in DMSO and BrdU CM from oligodendrocytes showing factors that are upregulated in BrdU CM treated (red) (log2fc > 1, p-value < 0.05) and downregulated in BrdU CM treated (blue) (log2fc < -1, p-value < 0.05). Black data points indicate factors that do not reach the threshold (−1 < log2fc < 1 | p-value > 0.05). E) Volcano plot of Nomic Bio Core Immune panel cytokines in DMSO and BrdU CM from neurons showing factors that are upregulated in BrdU CM treated (red) (log2fc > 1, p-value < 0.05) and downregulated in BrdU CM treated (blue) (log2fc < -1, p-value < 0.05). Black data points indicate factors that do not reach the threshold (−1 < log2fc < 1 | p-value > 0.05). F) Venn diagram showing the number of upregulated factors specifically in astrocyte BrdU CM (23) and microglia BrdU CM (69) as well as the commonly upregulated factors (16). G) Z-score heatmap showing selected Core Immune panel (Nomic Bio) cytokine expression in all cell types (endothelial cells (pink), microglia (yellow), oligodendrocytes (green), astrocytes (purple), and neurons (blue)) based on nELISA analysis of CM from DMSO (grey) and BrdU (red) treated cell lines (n=3 replicates). H) Normalized expression of CCL2 based on human cytokine array in DMSO CM (grey) and BrdU CM (red) from astrocytes, endothelial cells, and microglia (n=4 replicates). Normalized expression of MIF based on human cytokine array in DMSO CM (grey) and BrdU CM (red) from astrocytes, endothelial cells, and microglia (n=4 replicates). I) Normalized nELISA signal of CCL2 in DMSO CM (grey) and BrdU CM (red) from astrocytes, endothelial cells, microglia, oligodendrocytes, and neurons (n=3 replicates). Normalized nELISA signal of MIF in DMSO CM (grey) and BrdU CM (red) from astrocytes, endothelial cells, microglia, oligodendrocytes, and neurons (n=3 replicates). Data was analyzed by two-way ANOVA with Tukey’s multiple comparisons test (H) and two-way ANOVA with Šídák’s multiple comparisons test (I). All graphs show mean with error bars depicting standard deviation (ns, p>0.05, *** p<0.001).

Focusing specifically on the primary drivers of the spreading of senescence (astrocytes and microglia) we observed an upregulation of 23 cytokines in astrocyte BrdU CM and 69 in microglia BrdU CM, with 16 factors commonly upregulated in both (Figure 3F). To validate the specific SASP factors in astrocyte and microglia BrdU CM, we utilized a human cytokine array, which paneled 36 human cytokines (13 of which were also included in the nELISA list). Because this method is less sensitive than nELISA, it enables the detection of only the most highly expressed factors within each cell type’s senescent CM. We observed a significant upregulation of CCL2 specifically in astrocyte BrdU CM and a microglia-specific upregulation of MIF in their BrdU CM (Figure 3H), highlighting these two ligands as key cell-type-specific SASP factors which may contribute to these cell type’s ability to drive a secondary senescence phenotype in other brain cells. Interestingly, pairwise comparisons from our nELISA data showed that CCL2 was significantly upregulated in the BrdU CM of all cell types other than endothelial cells (Figure 3I), aligning strongly with the SA β-gal results demonstrating that senescent endothelial cells were the only cell type incapable of inducing secondary senescence in any other brain cell type (Figure 1G). Similarly, MIF expression was significantly upregulated in microglia, oligodendrocyte, and neuron BrdU CM, with a trending increase in astrocyte BrdU CM, but was significantly decreased in endothelial cell CM (Figure 3I). This expression pattern further supports our SA β-gal results (Figure 1G). Overall, these results highlight specific SASP factors in senescent astrocyte and microglia CM, specifically CCL2 and MIF, as potential mediators of the cell-type-specific spreading of senescence.

### Analysis of ligands and receptors in primary and secondary senescent cells

We next sought to investigate the opposing question of cell-type vulnerability to secondary senescence based on SASP receptor-ligand interactions. To do so, we applied the BulkSignalR algorithm (*29*) to our bulk RNAseq dataset from control and senescent astrocytes, endothelial cells, microglia, oligodendrocytes, and neurons (*13*) (Figure 4A). We isolated inferred activated ligand-receptor pairs based on altered downstream-regulated gene pathways in our senescent cell types (Figure 4B). Focusing on receptor expression, we identified both unique and common receptors across the various human brain cell types. Based on our SA β-gal results (Figure 1 D-G), which showed that endothelial cells, astrocytes, and microglia were undergoing secondary senescence in response to SASP, we prioritized receptors common to these cell types. This analysis identified three shared candidates (highlighted in red): CXCR7, KREMEN2, and GIPR (Figure 4B). Only one of these receptors, CXCR7, showed cell-type-specific expression restricted to the cell types that were capable of adopting a secondary senescence phenotype (astrocytes, endothelial cells, microglia) and was absent in the non-receivers (oligodendrocytes, neurons) (Figure 1G, 4C, S3B), making it a promising candidate for better understanding what makes these cells susceptible to adopting a secondary senescence phenotype. The CXCR7 receptor binds to the ligand CXCL12, which was significantly decreased in senescent (red) astrocytes but significantly increased in senescent (red) neurons (Figure 4C). We investigated expression levels of DPP4 in senescent (red) cells, which is known to cleave and inactivate CXCL12 (*23*), which was significantly upregulated in senescent astrocytes, endothelial cells, and neurons and showed a trending increase in senescent microglia (Figure 4C). Taken together, these data indicate a potential role of the CXCR7 receptor and DPP4 cleavage of CXCL12 in non-senescent receiving cells that are responsive to SASP, allowing for a spreading of senescence.

**Figure 4:**
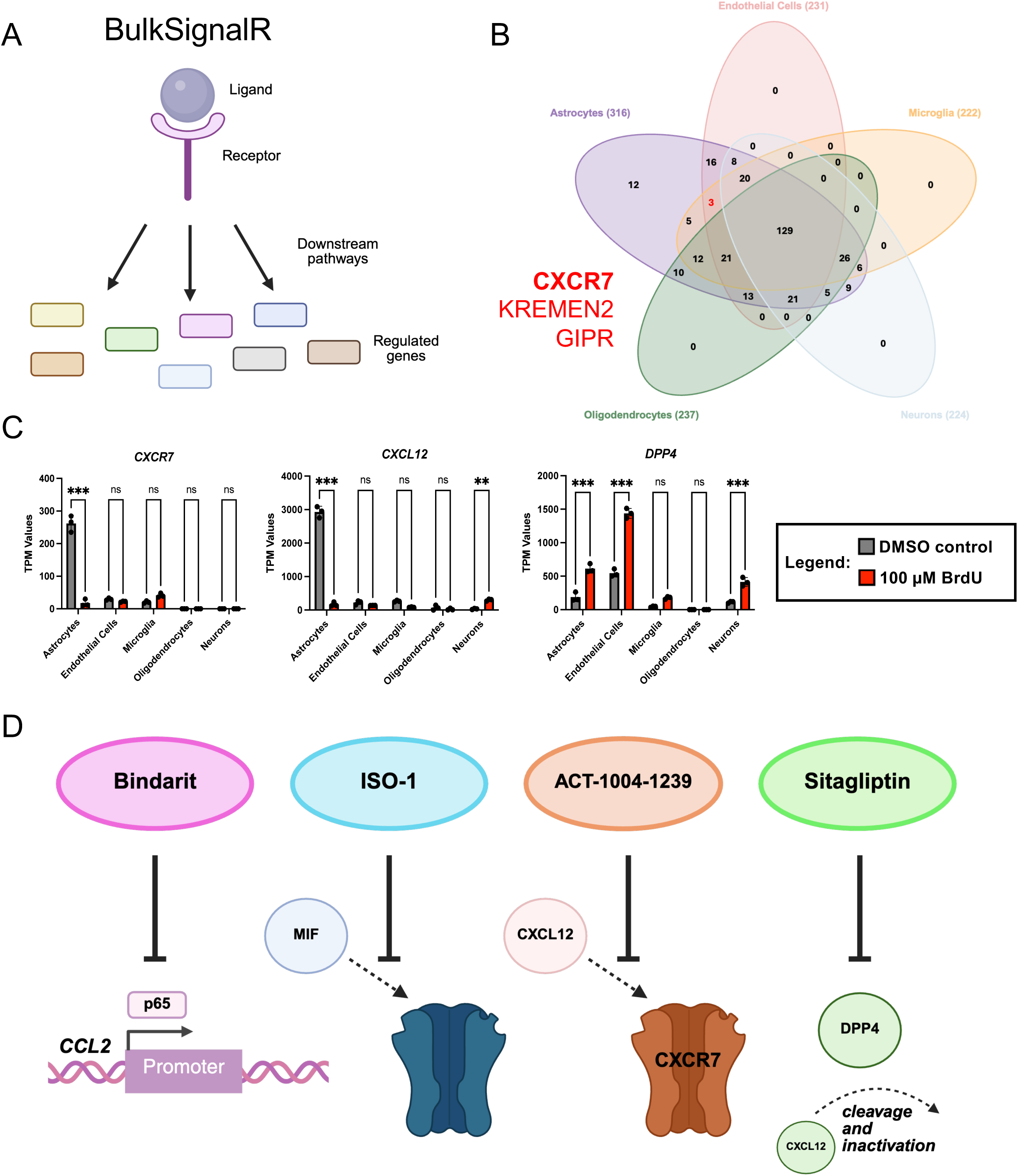
Analysis of ligands and receptors in senescent and receiving cells. A) Schematic depicting BulkSignalR pipeline which uses known ligand-receptor interactions and affected downstream pathways to analyze their activation based on our bulk RNAseq data from DMSO and BrdU treated human cell lines (created with BioRender). B) Venn diagram showing the number of receptors inferred from BulkSignalR to be activated across each of the five human cell types. Three receptors were identified in common between astrocytes (purple), endothelial cells (pink), and microglia (yellow) which were the cell types shown (Figure 1G) to be capable of receiving senescence signals and becoming SA β-gal positive: CXCR7, KREMEN2, and GIPR. Only CXCR7 was expressed in the cell types capable of entering secondary senescence (astrocytes, endothelial cells, microglia) (Figure 4C, S3B). C) TPM expression values of *CXCR7*, its ligand *CXCL12*, and *DPP4* which cleaves and inactivates CXCL12 in DMSO (grey) and BrdU (red) treated cell lines (n=3 replicates). D) Schematic of the four selected SASP inhibitors mechanisms of action: Bindarit is a CCL2 synthesis inhibitor which prevents p65 activation of the *CCL2* gene at the promoter region, ISO-1 is a MIF antagonist, ACT-1004-1239 is a CXCR7 antagonist, and Sitagliptin inhibits DPP4 preventing its action of cleaving and inactivating CXCL12 (created with BioRender). Data was analyzed by two-way ANOVA with Tukey’s multiple comparisons test (C). All graphs show mean with error bars depicting standard deviation (ns, p>0.05, ** p<0.01, *** p<0.001).

To directly test the role(s) of the identified ligands and receptors - including CCL2, MIF, CXCR7, and DPP4/CXCL12 - we selected four target-specific “SASP inhibitors” to use for treatments aiming to prevent a spreading of senescence between the human brain cell types (Figure 4D). Bindarit is a CCL2 synthesis inhibitor that prevents p65 activation of the *CCL2* gene at the promoter region and has been shown to be able to target brain cell types including astrocytes, endothelial cells, and microglia (*30*), which are the cell types we observed as capable of becoming senescent in response to SASP (Figure 1G). We confirmed in our bulk RNAseq data (*13*) that each of the five human brain cell types expressed at least one receptor for CCL2 (Figure S3A). ISO-1 is a MIF antagonist that has been harnessed as a pharmacological treatment for immune disorders (*31*). ACT-1004-1239 is a CXCR7 antagonist, which is a receptor that binds the ligand CXCL12, and Sitagliptin is a DPP4 inhibitor, which is the enzyme that cleaves and inactivates CXCL12 (*23*) (Figure 4D).

### Targeting SASP ligands and receptors to prevent the spreading of senescence

To determine the appropriate treatment concentrations for each of the four target-specific SASP inhibitors, astrocytes and microglia were treated with DMSO (grey) or BrdU (red) to induce senescence (*13*) along with varying concentrations of each drug (Bindarit – pink, ISO-1 – blue, ACT-1004-1239 – orange, and Sitagliptin – green) for 7 days (Figure S4). Based on our analysis, we selected the highest concentration of each drug that did not result in a significant decrease in cell viability independently for each cell type: 200 µM Bindarit (astrocytes) and 20 µM Bindarit (microglia), 50 µM ISO-1 (both astrocytes and microglia), 200 µM ACT-1004-1239 (astrocytes) and 2 µM ACT-1004-1239 (microglia), and 2 µM Sitagliptin (both astrocytes and microglia). We next determined if treatment with these SASP inhibitors for 7 days during 100 µM BrdU treatment altered the induction of a senescence phenotype, which could indicate a cell-autonomous spreading of senescence within the initial treatment plate as an autocrine-like propagation of senescence in neighboring cells of the same cell type following the initial treatment. After 7-day treatment with a 100 µM BrdU control (red) or 100 µM BrdU along with each of the four SASP inhibitors, we observed a significant reduction in the percentage of SA β-gal positive astrocytes with all drugs other than Bindarit (Figure 5A). Since Bindarit is a CCL2 synthesis inhibitor, it may not impact the induction of senescence since it relates to the production of what we believe is a key ligand, whereas the other SASP inhibitors are all related to the non-senescent receiver cells. In microglia, we observed a significant reduction in the induction of senescence only with co-treatment of 100 µM BrdU and the DPP4 inhibitor Sitagliptin (Figure 5B), which may indicate a specific role of CXCL12 in an autocrine-like spreading of senescence within this cell type.

**Figure 5:**
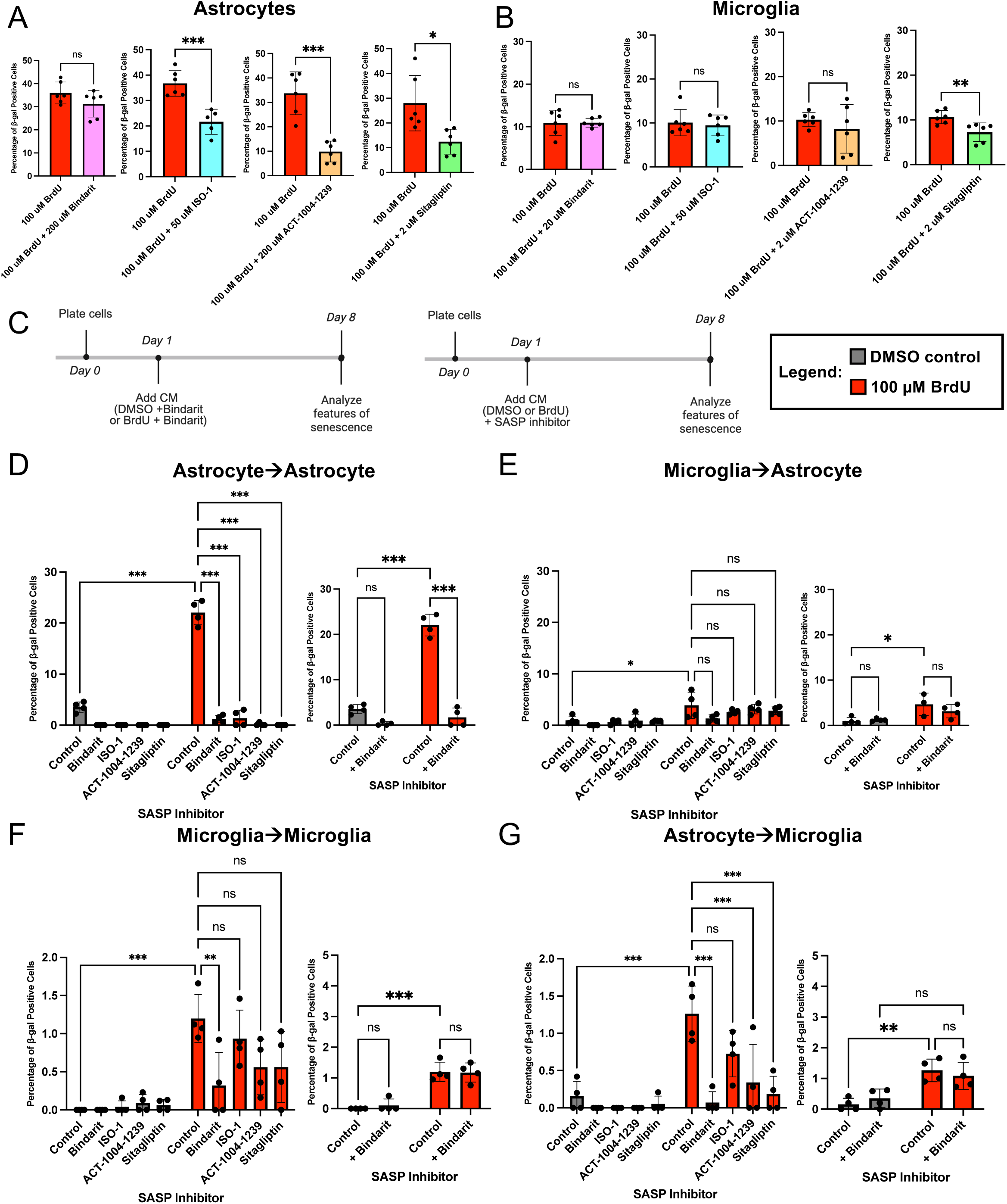
Targeting SASP ligands and receptors to prevent the spreading of senescence. A) Quantification of percentage of SA β-gal positive astrocytes following 7-day treatment with 100 µM BrdU (red) along with 200 µM Bindarit (pink), 50 µM ISO-1 (blue), 200 µM ACT-1004-1239 (orange), or 2 µM Sitagliptin (green) (n=6 replicates). B) Quantification of percentage of SA β-gal positive microglia following 7-day treatment with 100 µM BrdU (red) along with 200 µM Bindarit (pink), 50 µM ISO-1 (blue), 200 µM ACT-1004-1239 (orange), or 2 µM Sitagliptin (green) (n=6 replicates). C) Timeline showing treatment with DMSO + Bindarit CM or BrdU + Bindarit CM for 7 days. Timeline showing treatment with DMSO or 100 µM BrdU along with SASP inhibitors (ISO-1, ACT-1004-1239, or Sitagliptin) for 7 days. Features of senescence were analyzed 8 days after the initial plating of cells (created with BioRender). D) Quantification of percentage of SA β-gal positive astrocytes following 7-day treatment with DMSO + Bindarit CM from astrocytes (grey), BrdU + Bindarit CM from astrocytes (red), DMSO CM from astrocytes + SASP inhibitor (grey), or BrdU CM from astrocytes + SASP inhibitor (red) (n=4 replicates). Quantification of percentage of SA β-gal positive astrocytes following 7-day treatment with DMSO CM from astrocytes + Bindarit (grey) or BrdU CM from astrocytes + Bindarit (red) (n=4 replicates). E) Quantification of percentage of SA β-gal positive astrocytes following 7-day treatment with DMSO + Bindarit CM from microglia (grey), BrdU + Bindarit CM from microglia (red), DMSO CM from microglia + SASP inhibitor (grey), or BrdU CM from microglia + SASP inhibitor (red) (n=4 replicates). Quantification of percentage of SA β-gal positive astrocytes following 7-day treatment with DMSO CM from microglia + Bindarit (grey) or BrdU CM from microglia + Bindarit (red) (n=4 replicates). F) Quantification of percentage of SA β-gal positive microglia following 7-day treatment with DMSO + Bindarit CM from microglia (grey), BrdU + Bindarit CM from microglia (red), DMSO CM from microglia + SASP inhibitor (grey), or BrdU CM from microglia + SASP inhibitor (red) (n=4 replicates). Quantification of percentage of SA β-gal positive astrocytes following 7-day treatment with DMSO CM from microglia + Bindarit (grey) or BrdU CM from microglia + Bindarit (red) (n=4 replicates). G) Quantification of percentage of SA β-gal positive microglia following 7-day treatment with DMSO + Bindarit CM from astrocytes (grey), BrdU + Bindarit CM from astrocytes (red), DMSO CM from astrocytes + SASP inhibitor (grey), or BrdU CM from astrocytes + SASP inhibitor (red) (n=4 replicates). Quantification of percentage of SA β-gal positive astrocytes following 7-day treatment with DMSO CM from astrocytes + Bindarit (grey) or BrdU CM from astrocytes + Bindarit (red) (n=4 replicates). Data analyzed by unpaired t-test (A-B) and two-way ANOVA with Tukey’s or Šídák’s multiple comparisons test (D-G). All graphs show mean with error bars depicting standard deviation (ns, p>0.05, * p<0.05, ** p<0.01, *** p<0.001).

We went on to test treatment of the various SASP inhibitors alongside DMSO (grey) and BrdU (red) CM treatments to observe if we could prevent the paracrine spreading of senescence we previously observed between astrocytes and microglia (Figure 1D-G). Since Bindarit targets ligand synthesis, we generated CM from cells treated for 7 days with either DMSO + Bindarit or 100 µM BrdU + Bindarit to have a SASP composition containing all relevant factors other than CCL2. For CM treatments, we then exposed non-senescent cells to this DMSO/BrdU + Bindarit CM where we could assess if removing one specific SASP ligand would impact the spreading of senescence (Figure 5C). In the case of the other three SASP inhibitors, which focused more on impacting receptors on receiving non-senescent cells and which in one (or both) cell types influenced the initial induction of senescence during treatment (Figure 5A-B), we treated non-senescent cells with normal DMSO and BrdU CM from other cell types (collected as shown in Figure 1A) along with direct treatment of receiving cells with the various SASP inhibitors (Figure 5C).

When astrocytes were treated with DMSO (control, grey) and 100 µM BrdU (control, red) CM from other astrocytes, we recapitulated a significant increase in the percentage of SA β-gal positive senescent cells (Figure 1D, 5D). Interestingly, when treated with astrocyte BrdU + Bindarit CM and with astrocyte BrdU CM in the presence of ISO-1, ACT-1004-1239, or Sitagliptin, we were able to rescue the spreading of senescence between astrocytes (Figure 5D). We performed a control experiment with Bindarit where we used the alternate treatment type (Figure 5C) of normal DMSO or BrdU CM from astrocytes with Bindarit treatment directly on the non-senescent receiving cells, and we surprisingly observed a prevention of the spreading of senescence in this case as well (Figure 5D). This indicates that CCL2 may mediate an autocrine signal for the spreading of senescence between astrocytes but only in the context of CM-induced senescence. We went on to test the effects of the SASP inhibitor CM on astrocytes treated with CM from microglia. We similarly recapitulated the initial significant increase in the percentage of SA β-gal positive senescent cells we had observed (Figure 1D) but did not see any prevention of the spreading of senescence using any of the four SASP inhibitors in this case (Figure 5E). This indicates a cell-type specificity in the inducing cell’s SASP composition (independent of CCL2), regardless of the targeting of the non-senescent receiving cells. We performed the same Bindarit control experiment and did not observe a rescue of the significant upregulation of SA β-gal positive senescent astrocytes when treating microglia (Figure 5E).

When we utilized microglia as the receiving cell type, we assessed microglia-derived DMSO and BrdU CM treatment as a control, which showed the expected increase in the percentage of SA β-gal positive cells with BrdU CM (Figure 1F, 5F). Interestingly, treatment with BrdU + Bindarit CM from microglia was the only SASP inhibitor treatment that rescued this cell-autonomous spreading of senescence. The additional Bindarit control experiment in these cells also did not rescue the spreading of senescence (Figure 5F). Lastly, we examined microglia treated with CM from astrocytes. BrdU CM alone reiterated the increase in SA β-gal positive microglia (Figure 1F) and treatment with BrdU + Bindarit CM from astrocytes as well as with astrocyte BrdU CM along with the drugs targeting CXCR7 and DPP4 (ACT-1004-1239 and Sitagliptin, respectively) was able to prevent the spreading of senescence from astrocytes to microglia (Figure 5G). Finally, the Bindarit control experiment using DMSO or BrdU CM from astrocytes, along with Bindarit treated directly onto receiving microglia, was insufficient to rescue the spreading of senescence (Figure 5G). Overall, these results demonstrate that various SASP inhibitors can prevent the spreading of senescence in a cell-type-specific manner, highlighting their potential as therapeutic interventions to limit senescence propagation in disease contexts.

## Discussion

The work presented here highlights the cell-type-specific nature of the SASP, which mediates a paracrine spreading of senescence between various human brain cell types where there are key primary senescent drivers and secondary senescent receivers. Most importantly, we elucidated that astrocytes and microglia appear to be the main drivers of both cell-autonomous and cell non-autonomous paracrine spreading of senescence to other glial cells as well as endothelial cells. This important discovery informs models of neuron-glia interactions in neurodegenerative disease, suggesting that glial cells may drive of senescence, blood-brain barrier breakdown, inflammation, and ultimately contribute to neurodegenerative phenotypes. Additionally, we showed that it is possible to delineate key SASP ligand-receptor pairs that directly mediate this spreading phenotype. For example, we identified that CCL2, MIF, CXCR7, and DPP4-mediated cleavage of CXCL12 are critical factors for the spread of senescence in a glial cell-type-specific manner. When we directly targeted these ligands and receptors in astrocytes and microglia, we were able to in some cases prevent the paracrine spreading of senescence. Taken together, our data demonstrates a mechanism by which glial cells mediate the propagation of secondary senescence phenotypes in other brain cell types, potentially contributing to local brain inflammation and neuronal loss during aging and neurodegenerative disease.

nELISA profiling of conditioned media from DMSO- and BrdU-treated astrocyte, endothelial, microglial, oligodendrocyte, and neuronal human cell lines revealed cell-type-dependent upregulations of various cytokines involved in the SASP (Figure 1C). Although there are established characterizations of the SASP across multiple tissues (*32*), the work we present here offers a clear view into specific changes across specific brain cell types utilizing our well-characterized *in vitro* BrdU-induced senescence model (*13*). One major limitation is that we utilize human cancer cell lines to model senescence, which do not fully resemble differentiated cells *in vivo*, although they offer a highly controlled and cell-type-specific experimental approach. When we sought to elucidate which of the SASP upregulations may drive a paracrine spreading of senescence and in which cell types, we observed CM-induced SA β-gal expression in a cell-type-specific manner (Figure 1D-F). SA β-gal expression is an established biomarker of senescent cells, recently included in a list of minimal criteria for defining senescent cells *in vivo* (*33, 34*), which we have shown is characteristic of the human cell lines presented here following BrdU-induced senescence (*13*). However, there is debate in the field of cellular senescence regarding the low percentage of detectable SA β-gal positive cells across multiple models as well as the lack of specificity as a marker of ‘neurescent’ cells (*35, 36*). In line with this, we observed a relatively low percentage of SA β-gal positive cells in response to CM treatments, specifically when microglia were the receiving cell type.

Interestingly, it appeared that astrocytes and microglia were the main drivers of a paracrine spreading of senescence, where they acted as primary senescent cells that were able to induce secondary senescence in each other, in themselves, and in endothelial cells (Figure 1D, F). Alternatively, endothelial cells were able to enter a state of secondary senescence (by neuronal BrdU CM in addition to that from astrocytes and microglia), but their senescent CM was not sufficient to induce a paracrine spreading senescence in any other brain cell type (Figure 1E). Senescent oligodendrocytes and neurons could induce senescence in other cell types, but themselves were not able to adopt a ‘secondary senescence’ phenotype in response to BrdU CM treatment from any other cell type. Additionally, we demonstrated that the transmitted ‘secondary senescence’ phenotype is also cell-type-specific and that this differs from a fully penetrant, DNA damage-induced model of senescence in these same human brain cell lines (*13*) where it has a more subtle, nuanced expression of senescence hallmarks that is also dependent on the primary senescent cell type. Taken together, these results offer a unique cell-type-specific directionality, defining certain cell types as “senescence spreaders”, which are cells that can enter ‘primary senescence’ and induce ‘secondary senescence’ in others.

Our results further suggest a cell-type-specific susceptibility to senescence, distinguishing between secondary senescence receivers and non-receivers (Figure 1G). In the context of brain aging and neurodegenerative disease, our data suggest glial cell types as key mediators of SASP-induced senescence whereas endothelial cells of the blood-brain-barrier may adopt solely a ‘secondary senescence’ phenotype. This could lend to a model of local inflammation, where senescent glial cells could induce senescence in endothelial cells of the blood-brain-barrier, leading to blood-brain-barrier breakdown and loss of integrity resulting in CD8+ killer T-cell infiltration of the brain (*2*). Overall, our findings support our previous reports of cell-type-specificity across multiple senescence hallmarks in these brain cell types (*13*), while also showing evidence of a paracrine spreading of senescence that can be transmitted between different cell types (*19*). It is plausible that different cells display heterogeneous profiles of senescence (*12*) partially due to the molecular diversity between primary and secondary senescence which would be very difficult to delineate *in vivo*.

When we focused on the unique SASP factors from CM of key primary senescent drivers (astrocytes and microglia), we revealed that one astrocyte-specific cytokine (CCL2) and one microglia-specific cytokine (MIF) were the most strongly upregulated in senescent CM (Figure 3H). Interestingly, CCL2 has been previously described as a key SASP factor in mediating a paracrine spreading of senescence in fibroblasts (*18, 19*) and was targeted via inhibition of its receptor in bystander receiving cells. Further analysis revealed key receptors in the three cell types that we observed to be capable of adopting a ‘secondary senescence’ phenotype in response to senescent CM from at least one other cell type (Figure 4A, 1G). Only one of these identified receptors (CXCR7) was expressed selectively in the cell types capable of becoming senescent in response to the SASP of other cells (astrocytes, endothelial cells, and microglia) (Figure 4B, C, S3B). Additionally, although one of the ligands for the receptor (CXCL12) was generally decreased in the senescent CM of most cell types (Figure 3), the DPP4 enzyme was significantly upregulated in many BrdU-treated cell types which cleaves CXCL12 and has been previously associated with senescence (*23, 24*). In the context of atherosclerosis, it was shown that vascular smooth muscle cells (VSMCs) express DPP4 at elevated levels, which correlated with enhanced expression of SASP factors (*37*). Additionally, it was observed that blocking DPP4 had positive senolytic effects on VSMCs and the inflammatory phenotype, suggesting DPP4 as a target for senolytics and senomorphics. When we utilized SASP inhibitors to target these ligands and receptors, we revealed cell-type-dependent effects on the paracrine spreading of senescence.

Taken together, these results offer CCL2, MIF, CXCR7, and DPP4 as key mediators of a cell-type-specific paracrine spreading of senescence between glial cells and other human brain cell types. Ultimately, our findings identified astrocytes and microglia as key senescence inducers (spreaders) as well as specific ligands and receptors related to the SASP that mediate their ability to induce secondary senescence phenotypes in other brain cells. Future investigations can work to apply these findings in primary cells or in *in vivo* models to determine if these same cell-type-specific relationships mediate a spreading of senescence across various contexts. Ultimately, the work presented here informs the nature of a paracrine spreading of senescence between key human brain cell types and offers potential therapeutic targets for preventing this spread.

## Methods

### Culture of human cell lines

HMC3 cells (ATCC, Catalog No. CRL-3304), SK-N-MC cells (ATCC, Catalog No. HTB-10), HBEC-5i microvascular endothelial cells (ATCC, Catalog No. CRL-3245), HOG cells (Sigma, Catalog No. SCC163), and SVG-A astroglia cells (kindly provided by Dr. Walter Atwood, Brown University, Providence, RI, USA) were cultured as reported previously (*13*) and according to vendors’ protocols where applicable.

### BrdU treatment in cell culture

Treatment for 7 days with 100 µM 5-bromo-2’-deoxyuridine (BrdU) (Sigma, Catalog No. B5002) dissolved in DMSO (Sigma, Catalog No. D2650) was used to induce senescence as we have previously described (*13*). A 0.1% DMSO solution in the appropriate cell line growth media was used as a control.

### Collection of conditioned media (CM) and treatment in human cell lines

As described previously (*13*), to generate conditioned media (CM) from DMSO and BrdU treated cells, we treated the human cell lines for 7 days with DMSO and BrdU (as described in BrdU treatment in cell culture). Cells were then media changed and cultured for an additional 24 hours to condition the media (without the DMSO/BrdU treatment present). This CM was then collected, filtered, snap frozen, and stored at -80°C. For CM treatments, cells were given a 2:1 treatment of CM:normal media with a DMSO CM and BrdU CM group. Additionally, a media control group was added to isolate any effects of culturing one cell line in the media of another cell type. CM treatment was incubated on the cells for 7-days prior to assessment of senescence phenotypes.

### Senescence-associated β-galactosidase (SA β-gal) assay

To test whether cells displayed a senescence phenotype as we have reported previously (*13*), a senescence-associated β-galactosidase (SA β-gal) assay (Cell Signaling Technology, Catalog No. 9860) was performed according to the manufacturer’s protocol. After the respective 7-day treatment, cells were fixed and incubated over night at 37°C with X-gal as substrate for the SA β-galactosidase. The following day, the substrate solution was removed, 70% glycerol (in PBS) was added to the fixed cells, and wells were imaged with a microscope (Accu-Scope, EXI-410, Skye Color Camera). Analysis and quantification of images was performed using Fiji and statistical analyses were performed using Prism (GraphPad).

### RNA isolation and RT-qPCR

As we have previously reported (*13*), RNA was isolated from human cell lines using the RNeasy Plus Mini Kit (QIAGEN, Catalog No. 74134) and QIAshredder (QIAGEN, Catalog No. 79656). RNA (200 ng) was reverse transcribed using the Applied Biosystems High-Capacity cDNA Reverse Transcription Kit (Thermo Scientific, Catalog No. 4368814) and the 20 μL output was diluted in nuclease-free water for a cDNA working concentration of 5ng/μL. Real time PCR was performed using the Applied Biosystems *Power* SYBR Green PCR Master Mix (Thermo Scientific, Catalog No. 4367659) on a Quant Studio 3 (Applied Biosystems) with reaction specificity confirmed by melt curve analysis. All comparisons (ex. DMSO CM vs. BrdU CM) for each qPCR reaction were run on the same qPCR plate and run in triplicate. Human qPCR primer sequences used (5’ to 3’) are listed in Table 1.

**Table 1:**
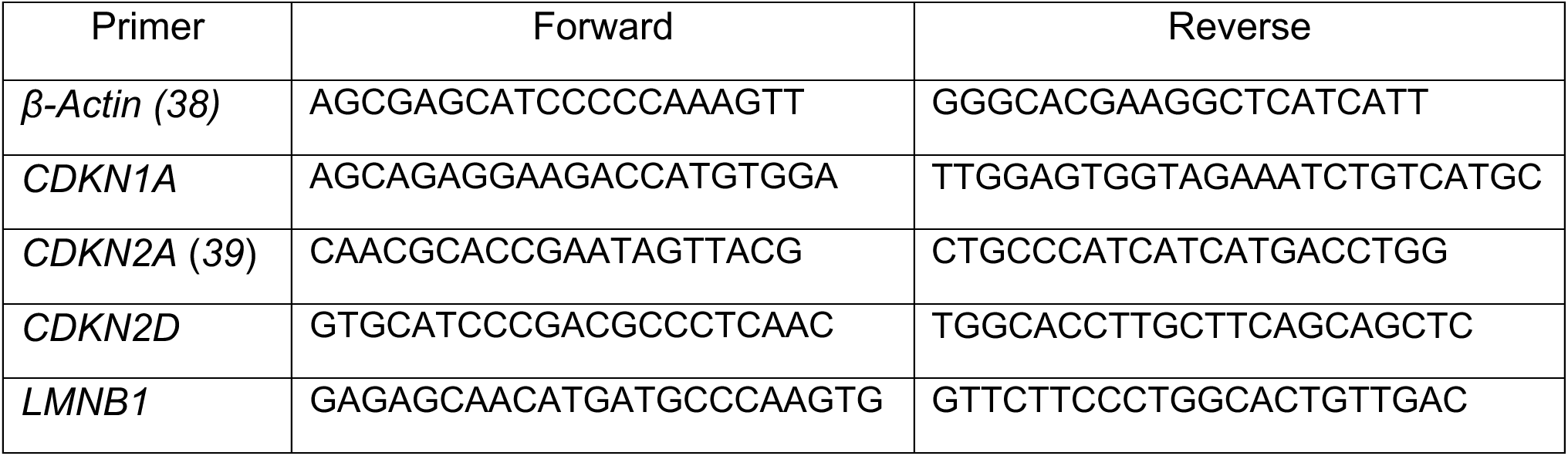
Human qPCR primers used to detect senescence hallmark expression.

### Immunofluorescent staining

As reported previously (*13*), 12 mm round glass coverslips (VWR, Catalog No. 73605-514) were washed in 70% ethanol, rinsed with PBS, and allowed to air dry for 60 minutes under UV light to sterilize. Coverslips were placed into each well of a 24-well plate, 500 µL of 0.1% gelatin in H_2_O was added to each well, and the plate was incubated at 37°C for at least 30 minutes. The gelatin solution was then removed, and cells were plated onto coverslips. Following treatment, cells were chemically fixed with 4% PFA in PBS for 20 minutes at room temperature. Cells were then permeabilized with 0.2% TritonX-100 in PBS for 10 minutes at room temperature followed by three 5-minute PBS washes and then blocking in 5% NDS/PBS/0.1% Tween20 at room temperature for 30 minutes. The cells were then incubated with the respective primary antibody (1:500) in 5% NDS/PBS/0.1% Tween20 overnight at 4°C. Primary antibodies used include γH2AX (pS139) (Abcam, Catalog No. ab303656, Lot 1099105-4, Clone N1-431), lamin B1 (Abcam, Catalog No. ab16048, Lot 1085092-1), and p21 (Abcam, Catalog No. ab109520, Lot 1092672-16). The following day, cells were washed 4 times for 5 minutes each in 0.1% Tween20-PBS and then incubated with the respective secondary antibodies (1:1000) in 5% NDS/PBS/0.1% Tween20 for 2 hours in the dark at room temperature. Secondary antibodies used include Alexa Fluor 488 d@ms IgG (Invitrogen, Catalog No. A21202, Lot 3006753) and Alexa Fluor 488 d@Rb (Invitrogen, Catalog No. A21206, Lot 2668665). The cells were then rinsed again 4 times for 5 minutes each in 0.1% Tween20-PBS and then coverslips with cells were mounted onto slides (top down) with 20 uL of DAPI Fluoromount-G (Southern Biotech, Catalog No. 0100-20). Slides were then cured in the dark at room temperature overnight. Confocal images were taken with an Olympus FV3000 Laser Scanning Confocal Microscope. Laser settings (laser strength, gain, and offset) and magnification were maintained across treatment groups.

### Fluorescent dyes for imaging mitochondria and lysosomes

For imaging of mitochondria and lysosomes as we have described previously (*13*), cells were washed 1x with Dulbecco’s PBS without calcium and magnesium (Thermo Scientific, Catalog No. 14190094) and then incubated for 15 minutes at 37°C in either 500 nM Mitotracker Red CMXRos (Invitrogen, Catalog No. M7512) in PBS or 1 µM Lysotracker Deep Red (Invitrogen, Catalog No. L12492) in PBS. Following treatment, cells were chemically fixed, permeabilized, washed, blocked, and stained as described in Immunofluorescent staining for additional senescence markers. Confocal images were taken with an Olympus FV3000 Laser Scanning Confocal Microscope. Laser settings (laser strength, gain, and offset) and magnification were maintained across treatment groups. Post-processing of images was performed by ImageJ and Cell Profiler as described in Image analyses.

### Image analyses

All Olympus FV3000 images were collected at 60x magnification, 1024x1024 resolution, and with optimal Z-stacks as reported previously (*13*). Images were analyzed in bulk through Cell profiler. Z-projections were taken from each image by maximum intensity and then separated by fluorophore. Nuclei were identified using DAPI staining and cell types were identified through specific cell markers. Mitochondria and lysosomal counts were analyzed by applying a size and intensity threshold in Cell Profiler utilizing control samples. Counts per image were then standardized by the number of cells to give an average number of objects per cell. γH2AX foci were analyzed by applying a size and intensity threshold in Cell Profiler utilizing control samples. These foci were then associated with each nucleus. Foci per nuclei were then averaged by the number of cells per image. p21 and lamin B1 intensity were extracted from DAPI labeled nuclei.

### Human cytokine array

A Proteome Profiler Array, Human Cytokine Array (R&D Systems, Catalog No. ARY005B) was performed according to the manufacturer’s protocol. Membranes were blocked in Array Buffer for 1 hour at room temperature. During this time, each CM sample (500 µL) was added to 1 mL of Array Buffer in separate tubes followed by the addition of 15 µL of Detection Antibody Cocktail to each sample, which was mixed and incubated for 1 hour at room temperature. Array Buffer was aspirated from the wells containing membranes and sample/antibody mixtures were added and incubated overnight at 4°C. The following day, membranes were removed and washed three times for 10-minutes each with 1X Wash Buffer. Diluted Streptavidin-HRP was added to each membrane and incubated for 30 minutes at room temperature. Membranes were then washed again three times for 10-minutes each with 1X Wash Buffer. Using forceps, membranes were carefully placed in plastic sheet protectors. Equal volumes of Chemi Reagents 1 and 2 were mixed and 1 mL of this mixture was added onto each membrane before covering the top layer of the plastic protector and incubating for 1 minute. Excess mix was then squeezed out of the plastic protector and a Kimwipe was used directly on the membrane to blot any remaining mixture. A ChemiDoc XRS+ (BioRad) was used to image membranes to visualize positive signals, which were subsequently analyzed using the overlay map of the 36 human cytokine positions. Each signal was an average from the duplicate spots of each cytokine followed by subtraction of average background signal from negative control spots on each membrane.

### NELISA (Nomic Bio)

Human cell line CM samples (50 µL each) were prepared and plated on a 96-well plate which was sealed, packaged, and sent to Nomic Bio for nELISA analysis using their Core Immune panel. Their nELISA package included sample processing and data upload to an online portal, which included raw signal (pg/mL), fluorescence intensity values, as well as normalized nELISA signals of 286 factors which we used to generate volcano plots and bar graphs comparing alterations in levels of various immune factors included in their panel. Samples used in heatmaps were normalized by z-score.

### Bulk Signal R

Ligand-receptor interactions between cell types were inferred using *BulkSignalR* (v1.0.2) (*29*). RNAseq count data from purified cell populations were combined and processed to create a *BSRDataModel* object with upper quartile (UQ) normalization. Statistical model parameters were learned using the *LearnParameters* function with quick mode enabled. To infer ligand-receptor interactions, we applied the *BSRInference* package and adjusted the minimum correlation threshold to 0.3 and the REACTOME pathway reference database. Significant interactions were identified using a q-value threshold of 0.01. Gene signature scores were calculated using *scoreLRGeneSignatures* to assess ligand-receptor interaction activity across samples. The direction of intercellular communication was determined by comparing mean ligand expression in the source cell type with mean receptor expression in the target cell type.

### RNAseq data from human cell lines

RNA sequencing of DMSO- and BrdU-treated human cell lines was performed and described previously (*13*). Here, we utilized TPM (transcripts per million) values to generate bar graphs for relevant transcripts.

### CCK8 Viability assay

To assess cellular viability, a CCK8 cell viability kit (Dojindo, Catalog No. CK04-05) was used according to the manufacturer’s protocol. As described previously (*13*), cell medium was replaced by fresh medium containing 10% of CCK8 solution in the appropriate serum-containing media based on the cell type being tested. Using a SpectraMax iD3 (Molecular Devices) plate reader, an initial background absorbance reading at 450 nm was taken. Cells were then incubated for 2 hours at 37°C with 5% CO_2_ before absorbance was read again.

### BrdU and SASP inhibitor treatment

To observe if treatment with SASP inhibitors (Bindarit, ISO-1, ACT-1004-1239, and Sitagliptin) alongside BrdU treatment altered the induction of senescence, we treated astrocytes and microglia for 7 days with each inhibitor along with 100 µM BrdU (as compared to a 100 µM BrdU control) and then performed a SA β-gal assay (as described in Senescence-associated β-galactosidase (SA β-gal) assay). This allowed for comparisons between the percentage of senescent cells with BrdU alone and with co-treatment of BrdU and each individual SASP inhibitor in both cell types.

### Bindarit CM collection

Treatment with the CCL2 synthesis inhibitor Bindarit (MedChemExpress, Catalog No. HY-B0498) (*30*) was used to specifically target CCL2 production in senescent cells. Bindarit was dissolved in DMSO to a stock concentration of 100 mM and was diluted in appropriate cell culture media. Following a CCK8 viability assay, 200 µM (astrocytes) and 20 µM (microglia) were selected as treatment conditions. Cells were simultaneously treated with 100 µM BrdU or DMSO control to induce senescence along with their respective concentration of Bindarit for 7 days. Cells were then media changed and cultured for an additional 24 hours to condition the media (without the DMSO/BrdU and Bindarit treatment present). This Bindarit CM was then collected, filtered, snap frozen, and stored at -80°C.

### Bindarit treatment

Non-senescent astrocytes and microglia were exposed to CM from DMSO or BrdU treated astrocytes and microglia for 7 days. This was compared to non-senescent astrocytes and microglia that were exposed to CM from DMSO + Bindarit or BrdU + Bindarit astrocytes and microglia for 7 days. Comparisons were made based on the percentage of senescent cells utilizing a SA β-gal assay (as described in Senescence-associated β-galactosidase (SA β-gal) assay). These experiments were set to determine the effect of removing CCL2 specifically from the senescent CM of astrocytes and microglia on their ability to induce senescence in non-senescent astrocytes and microglia.

### ISO-1 treatment

Treatment with migration inhibitor factor (MIF) antagonist ISO-1 (MedChemExpress, Catalog No. HY-16692) (*31*) was utilized to prevent MIF-associated communication between senescence inducers and receiving cells. ISO-1 was dissolved in DMSO to a stock concentration of 100 mM and was diluted in appropriate cell culture media. Following a CCK8 viability assay, 50 µM (both astrocytes and microglia) was selected as the treatment condition. During 7-day treatment with DMSO and BrdU CM from astrocytes and microglia, non-senescent astrocytes and microglia were also treated with 50 µM ISO-1. These groups were compared to non-senescent astrocytes and microglia that were treated with DMSO and BrdU CM from astrocytes and microglia alone. Cells from all groups were analyzed using a SA-β-gal assay (as described in Senescence-associated β-galactosidase (SA-β-gal) assay) to determine the percentage of senescent cells in each condition. These experiments were set to determine the effect of preventing MIF-dependent association with non-senescent receiver cells as a potential means to mitigate a spreading of senescence.

### ACT-1004-1239 treatment

Treatment with CXCR7 antagonist ACT-1004-1239 (MedChemExpress, Catalog No. HY-142617) was utilized to prevent CXCR7-associated communication between senescence inducers and receiving cells. ACT-1004-1239 was dissolved in DMSO to a stock concentration of 10 mM and was diluted in appropriate cell culture media. Following a CCK8 viability assay, 200 µM (astrocytes) and 2 µM (microglia) were selected as treatment conditions. During 7-day treatment with DMSO and BrdU CM from astrocytes and microglia, non-senescent astrocytes and microglia were also treated with their respective concentration of ACT-1004-1239. These groups were compared to non-senescent astrocytes and microglia that were treated with DMSO and BrdU CM from astrocytes and microglia alone. Cells from all groups were analyzed using a SA β-gal assay (as described in Senescence-associated β-galactosidase (SA β-gal) assay) to determine the percentage of senescent cells in each condition. These experiments were set to determine the effect of preventing CXCR7-dependent association with non-senescent receiver cells as a potential means to mitigate a spreading of senescence.

### Sitagliptin treatment

Treatment with DPP4 inhibitor Sitagliptin (MedChemExpress, Catalog No. HY-13749) was used to eliminate DPP4-dependent cleavage and inactivation of CXCL12, which is a ligand of the CXCR7 receptor. Sitagliptin was dissolved in DMSO to a stock concentration of 100 mM and was diluted in appropriate cell culture media. Following a CCK8 viability assay, 2 µM (both astrocytes and microglia) was selected as the treatment condition. During 7-day treatment with DMSO and BrdU CM from astrocytes and microglia, non-senescent astrocytes and microglia were also treated with 2 µM Sitagliptin. These groups were compared to non-senescent astrocytes and microglia that were treated with DMSO and BrdU CM from astrocytes and microglia alone. Cells from all groups were analyzed using a SA β-gal assay (as described in Senescence-associated β-galactosidase (SA β-gal) assay) to determine the percentage of senescent cells in each condition. These experiments were set to determine the effect of preventing DPP4 from cleaving and inactivating CXCL12 in non-senescent receiving cells as a potential means to mitigate a spreading of senescence.

### Bindarit control treatment

Treatment with Bindarit on non-senescent receiver cells was used as a control to examine the effect of blocking CCL2 synthesis during CM-induced induction of senescence. diluted in appropriate cell culture media. During 7-day treatment with DMSO and BrdU CM from astrocytes and microglia, non-senescent astrocytes and microglia were also treated with the same treatment conditions for each cell type based on CCK8 viability (200 µM (astrocytes) and 20 µM (microglia). These groups were compared to non-senescent astrocytes and microglia that were treated with DMSO and BrdU CM from astrocytes and microglia alone. Cells from all groups were analyzed using a SA β-gal assay (as described in Senescence-associated β-galactosidase (SA β-gal) assay) to determine the percentage of senescent cells in each condition.

### Statistics

Data was analyzed using Fiji (v2.9.0), R (v4.4.3), Cell Profiler (v4.2.8), GraphPad Prism (v10.6.1), and R studio (v026.01.0+392). Statistical tests were performed using GraphPad’s Prism software or R studio when samples sizes exceeded Prism’s data limits. A threshold of p<0.05 was considered significant. Significance was determined using the test indicated in figure legends. Results are presented as means ± SD, with individual points representing biological replicates. All sample sizes (n) are listed as either replicates, indicating individual cell culture wells. All measurements were taken from distinct samples.

---

Figures S1-S2 are associated with Figure 2. Figure S3 is associated with Figure 4. Figure S4 is associated with Figure 5.

**Figure S1.**
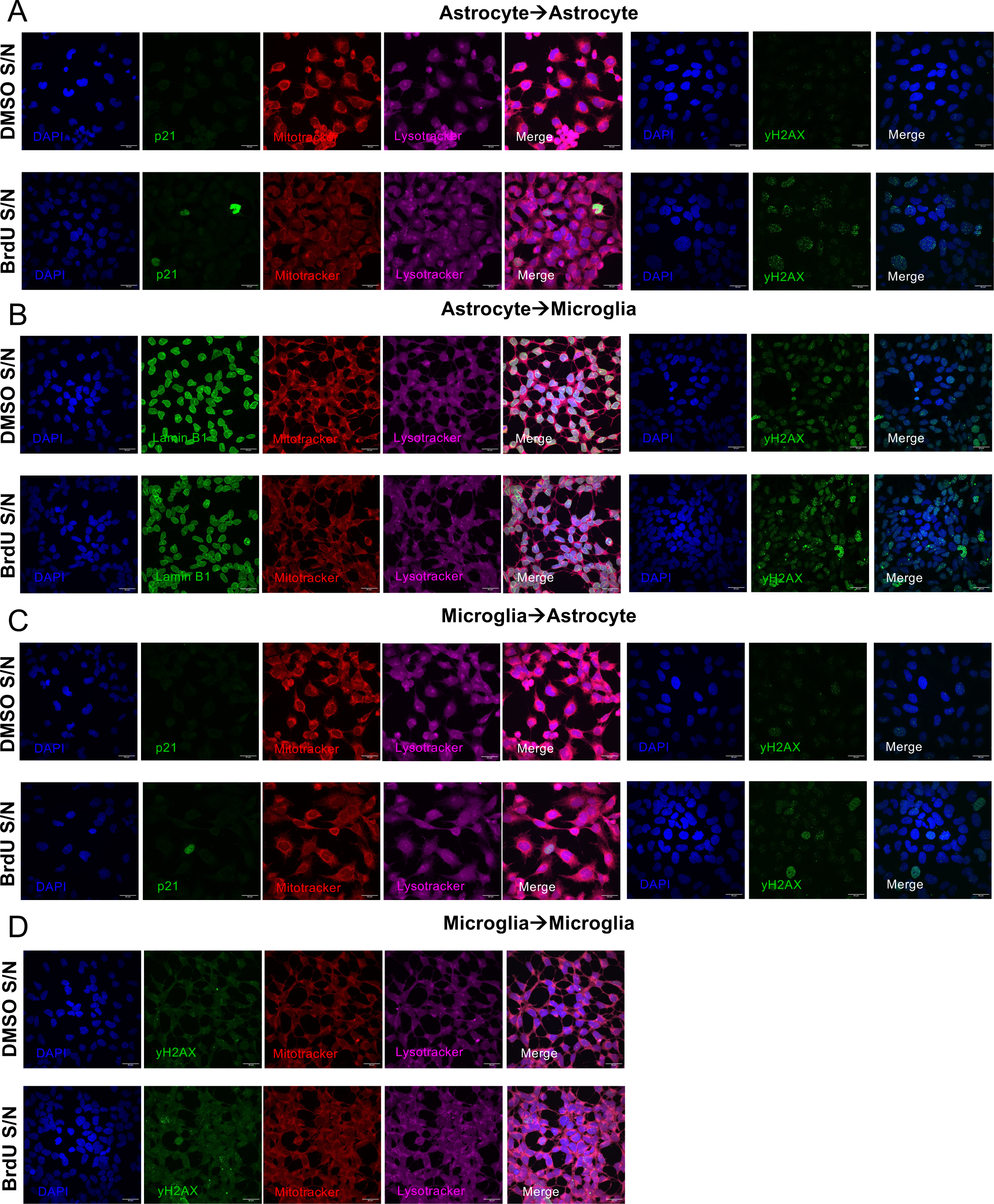
A) Confocal images of astrocytes treated with DMSO CM and BrdU CM from astrocytes with staining for p21 (green), Mitotracker Red CMXRos (red), Lysotracker (purple) and yH2AX (green) with DAPI (blue). B) Confocal images of microglia treated with DMSO CM and BrdU CM from astrocytes with staining for p21 (green), Mitotracker Red CMXRos (red), Lysotracker (purple) and yH2AX (green) with DAPI (blue). C) Confocal images of astrocytes treated with DMSO CM and BrdU CM from microglia with staining for p21 (green), Mitotracker Red CMXRos (red), Lysotracker (purple) and yH2AX (green) with DAPI (blue). D) Confocal images of microglia treated with DMSO CM and BrdU CM from microglia with staining for p21 (green), Mitotracker Red CMXRos (red), Lysotracker (purple) with DAPI (blue). All scale bars are 30 µm. *Note: a rendering error in the original preprint version caused a portion of the yH2AX DMSO S/N panel in A to display as partially blank black space, which has been corrected here to restore the complete original image data.*

**Figure S2.**
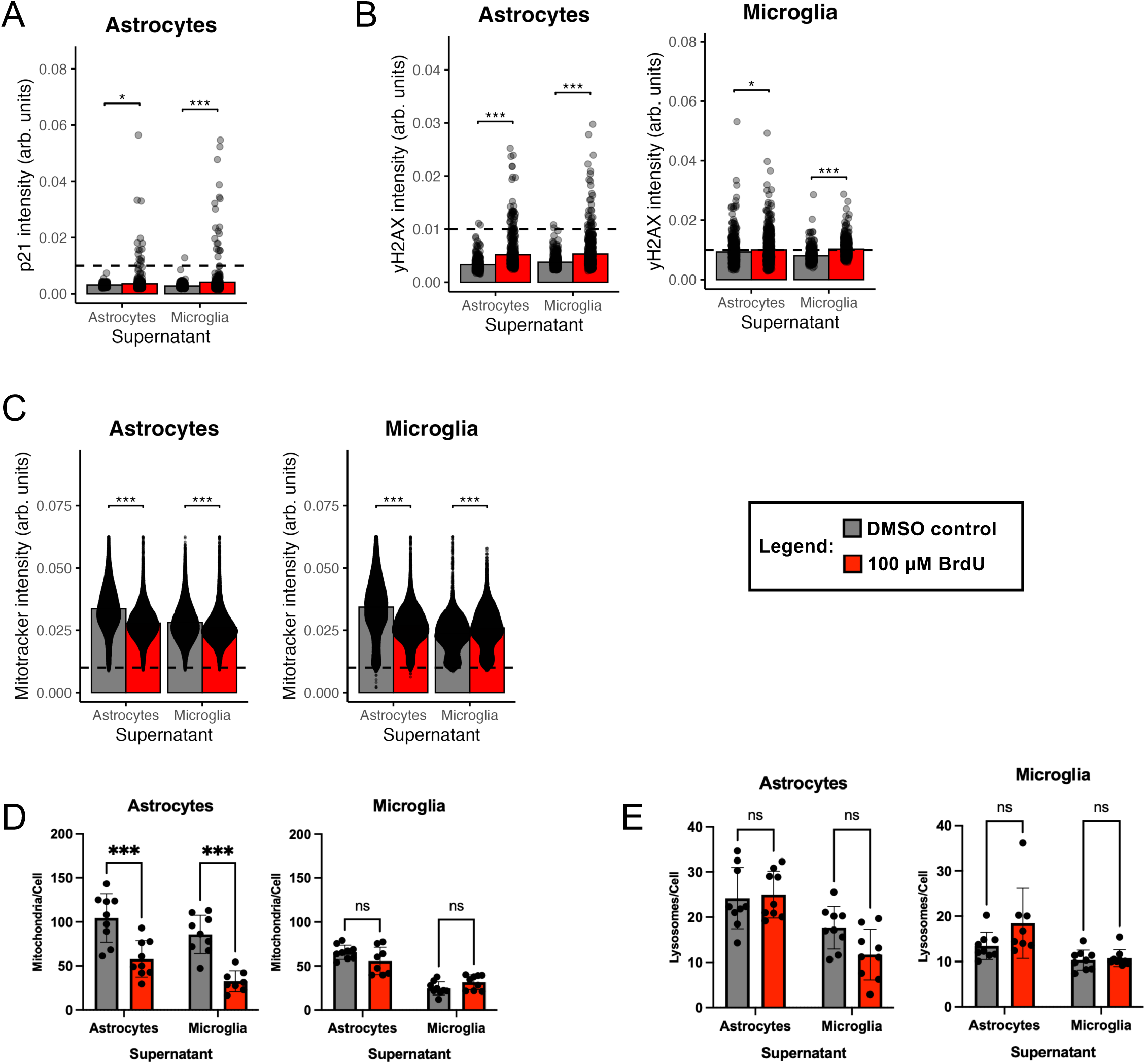
A) Quantification of p21 intensity in astrocytes treated with DMSO CM (grey) and BrdU CM (red) from astrocytes and microglia (n=9 replicates). B) Quantification of γH2AX intensity in astrocytes treated with DMSO CM (grey) and BrdU CM (red) from astrocytes and microglia (n=9 replicates). Quantification of γH2AX intensity in microglia treated with DMSO CM (grey) and BrdU CM (red) from astrocytes and microglia (n=9 replicates). C) Quantification of Mitotracker Red CMXRos intensity in astrocytes treated with DMSO CM (grey) and BrdU CM (red) from astrocytes and microglia (n=9 replicates). Quantification of Mitotracker Red CMXRos intensity in microglia treated with DMSO CM (grey) and BrdU CM (red) from astrocytes and microglia (n=9 replicates). D) Quantification of mitochondrial mass (mitochondria/cell) based on Mitotracker Red CMXRos staining in astrocytes treated with DMSO CM (grey) and BrdU CM (red) from astrocytes and microglia (n=9 replicates). Quantification of mitochondrial mass (mitochondria/cell) based on Mitotracker Red CMXRos staining in microglia treated with DMSO CM (grey) and BrdU CM (red) from astrocytes and microglia (n=9 replicates). E) Quantification of lysosomal mass (lysosomes/cell) based on Lysotracker staining in astrocytes treated with DMSO CM (grey) and BrdU CM (red) from astrocytes and microglia (n=9 replicates). Quantification of lysosomal mass (lysosomes/cell) based on Lysotracker staining in microglia treated with DMSO CM (grey) and BrdU CM (red) from astrocytes and microglia (n=9 replicates). Data was analyzed by two-way ANOVA with Tukey’s multiple comparisons test (A-E). All graphs show mean with error bars depicting standard deviation (ns, p>0.05, * p<0.05, *** p<0.001).

**Figure S3.**
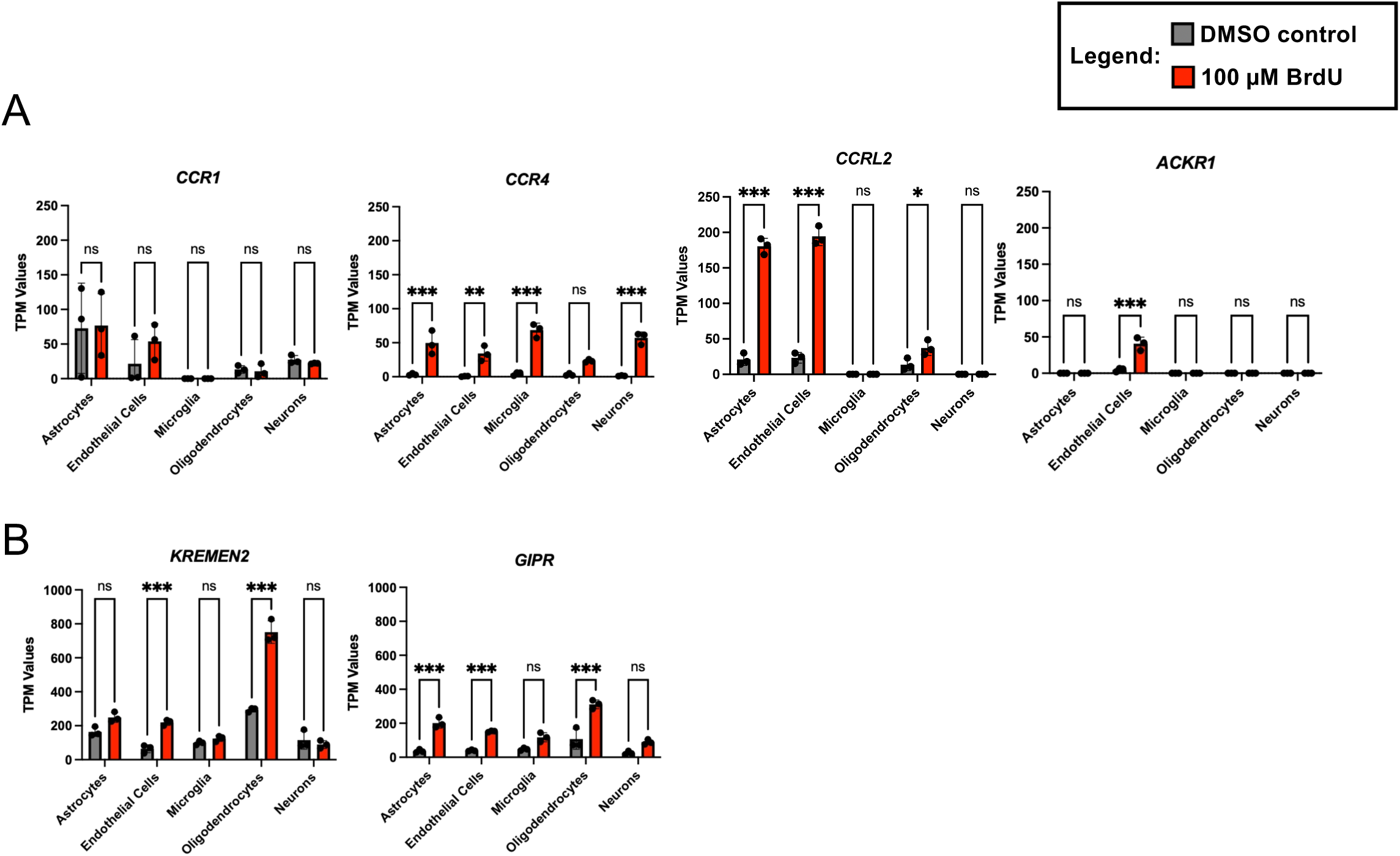
A) TPM expression values of CCL2 receptors *CCR1, CCR4, CCRL2,* and *ACKR1* in DMSO (grey) and BrdU (red) treated cell lines (n=3 replicates). B) TPM expression values of receptors *KREMEN2* and GIPR in DMSO (grey) and BrdU (red) treated cell lines (n=3 replicates). Data was analyzed by two-way ANOVA with Tukey’s multiple comparisons test (A-B). All graphs show mean with error bars depicting standard deviation (ns, p>0.05, * p<0.05, ** p<0.01, *** p<0.001).

**Figure S4.**
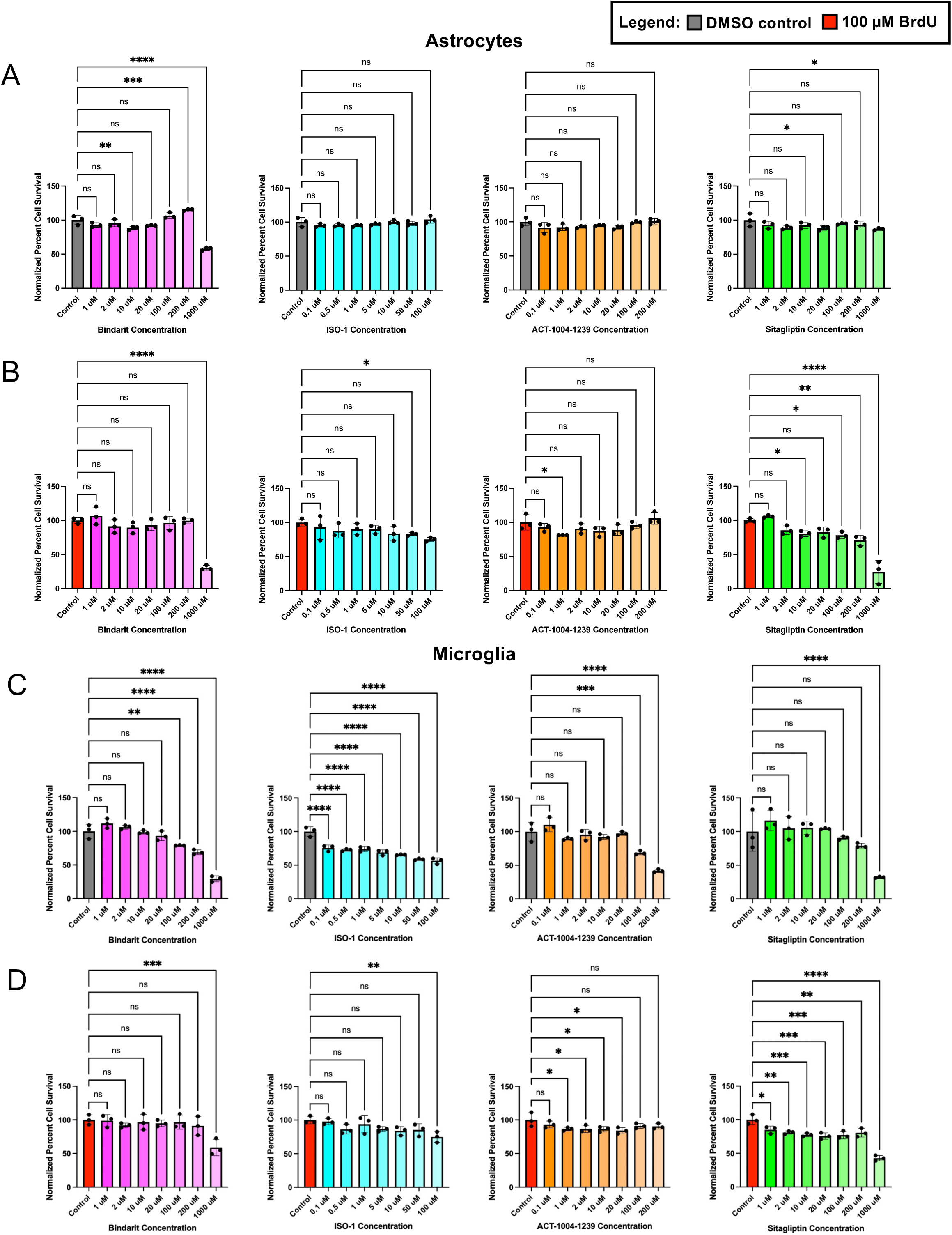
A) CCK8 viability assay in astrocytes showing normalized percent cell survival following treatment with DMSO control (grey) or DMSO + Bindarit (pink), DMSO + ISO-1 (blue), DMSO + ACT-1004-1239 (orange), or DMSO + Sitagliptin (green) at various concentrations (n=3 replicates). B) CCK8 viability assay in astrocytes showing normalized percent cell survival following treatment with 100 µM BrdU (red) or BrdU + Bindarit (pink), BrdU + ISO-1 (blue), BrdU + ACT-1004-1239 (orange), or BrdU + Sitagliptin (green) at various concentrations (n=3 replicates). C) CCK8 viability assay in microglia showing normalized percent cell survival following treatment with DMSO control (grey) or DMSO + Bindarit (pink), DMSO + ISO-1 (blue), DMSO + ACT-1004-1239 (orange), or DMSO + Sitagliptin (green) at various concentrations (n=3 replicates). D) CCK8 viability assay in microglia showing normalized percent cell survival following treatment with 100 µM BrdU (red) or BrdU + Bindarit (pink), BrdU + ISO-1 (blue), BrdU + ACT-1004-1239 (orange), or BrdU + Sitagliptin (green) at various concentrations (n=3 replicates). Data analyzed by one-way ANOVA with Dunnett’s correction for multiple comparisons (A-D). All graphs show mean with error bars depicting standard deviation (ns, p>0.05, * p<0.05, ** p<0.01, *** p<0.001).

## Acknowledgements and Funding

This work was supported in part through grants NINDS 1R01NS124735, the Hartman Foundation, and the Center for Healthy Aging, Stony Brook University. We wish to thank Dr. Jonathan Plessis-Belair for experimental design advice. We thank Wendy Akmentin for her invaluable help on all aspects of running lab facilities.

## Author Contributions

Conceptualization: TR, MR; Methodology: TR, MR; Investigation: TR; Visualization: TR; Funding acquisition: MR; Project administration: MR; Supervision: MR; Writing – original draft: TR; Writing – review & editing: TR, MR.

